# Nerve growth factor receptor identifies a basal subpopulation linked to poor prognosis and reduced immunotherapy responses in bladder cancer

**DOI:** 10.64898/2026.05.14.725085

**Authors:** Juan García-Agulló, Vanesa Santos, Mark Kalisz, Miriam Marqués, Elena Andrada, Catalina Berca, María Ramal, Jaime Martínez de Villarreal, Manuel Pérez-Martínez, Markus Eckstein, Raquel Benítez, Eduardo Caleiras, Núria Malats, Francisco X Real, Héctor Peinado

## Abstract

**Purpose:** Bladder cancer (BLCA) is a heterogeneous tumor type. Only one third of muscle-invasive (MIBC) patients respond to immune checkpoint inhibitors (ICIs). Reliable resistance markers are needed to guide clinical decisions. We investigated the nerve growth factor receptor (NGFR) in BLCA and analyzed its correlation with disease progression and response to immunotherapy.

**Experimental Design:** We analyzed NGFR expression in BLCA cell lines, organoids, mouse models and patient samples. The cohorts used were The Cancer Genome Atlas (TCGA), enriched in muscle-invasive bladder cancer (MIBC) (n=407); IMvigor210, representing MIBC patients treated with ICIs (n=348); and UROMOL2, as a non-muscle-invasive bladder cancer (NMIBC)-specific cohort (n=535). IMvigor010 was also included (n=728). Patients were stratified by NGFR expression quartiles. We analyzed survival and tumor subtypes and performed stromal deconvolution and functional profiling. We assessed stemness- and invasion-related features in SCaBER cells.

**Results:** NGFR marks a basal tumor cell subcluster and is independently associated with poor prognosis in TCGA and IMvigor210. NGFR-high tumors show stromal content enriched in cancer-associated fibroblasts, lower neoantigen burden, higher CD8^+^ T effector signature together with an immune-excluded phenotype, and a CAF-specific TGFβ signature. In the immunotherapy-treated cohort, high NGFR expression was also associated with poorer outcome. Functionally, NGFR appears to promote a stem-like/pro-invasive program in BLCA cells.

**Conclusions:** NGFR identifies a basal-like BLCA subpopulation linked to poor survival, while its association with immunotherapy response requires further validation. In addition, our in vitro analyses support a role of NGFR in stem-like and invasive traits, highlighting its relevance as a biomarker in BLCA.

## Introduction

Bladder cancer (BLCA) is a highly heterogeneous disease, classified into non-muscle invasive (NMIBC) and muscle-invasive subtypes (MIBC) [1]. Most diagnosed lesions are NMIBC, often having a good prognosis. In contrast, MIBCs comprise approximately 20% tumors that frequently metastasize to distant organs, including the liver, lung, and bone and are associated with high mortality [2]. Although MIBCs can arise from the progression of NMIBC, most are diagnosed *de novo* and present distinct molecular characteristics [1].

The molecular classifications of BLCA have focused on both NMIBC [3, 4] and MIBC [5, 6], distinguishing between luminal and basal subtypes. In NMIBC, three classes were proposed [3]. Class 1 lesions are low-grade luminal-like tumors with good prognosis. In contrast, Class 2 tumors present a worse prognosis and higher risk to progress to MIBC. Class 2 tumors, further divided into 2a (proliferative) and 2b (stem-like/EMT, stroma-rich) have worse prognosis [4]. Class 3 tumors have an intermediate prognosis. In MIBC, luminal and basal subtypes have been taxonomically distinguished based on gene expression [5], providing the foundation for the consensus classification currently used [6]. This classification closely parallels that of breast cancer, with basal tumors marked by worse outcomes, stem-like traits and cancer stem cell (CSCs) markers.

The origin of BLCA is still under debate, although it is proposed that urothelial basal cells play a fundamental role in the development of MIBC basal tumors [7, 8]. In a transgenic mouse model, Stat3 overexpression in basal cells induced MIBC directly upon carcinogen administration, bypassing NMIBC [7]. Basal urothelial cells coexpress SHH and KRT5 [9], with KRT14 identifying the most progenitor population [8]. In cancer, the expression of these markers is also apparent. KRT14^+^ cancer cells exhibit tumor-initiating capacity *in vivo* and are thus recognized as CSCs [8]. CSCs are a specialized subpopulation of tumor cells with stem-like properties that drive tumor growth and are key contributors to tumor recurrence and metastasis [10]. In BLCA, CSCs are a distinct subpopulation exhibiting self-renewal, tumorigenicity, and resistance to conventional treatments, playing a critical role in tumor initiation, progression, metastasis, and recurrence [11–13].

Immunotherapy has revolutionized cancer treatment [14]. In the context of BLCA, especially in MIBC tumors, immune checkpoint inhibitors (ICIs), such as pembrolizumab and atezolizumab, have become a major component of the therapeutic approach [14]. However, primary resistance to ICIs remains a significant challenge. Resistant tumors often exhibit characteristics typical of CSCs [15], proposing immune evasion as a newly recognized hallmark of CSCs [16]. In contrast to chemotherapy resistance, where the mechanisms associated with CSCs are better characterized, the effects of CSCs on the immune system and their role in ICI resistance remain less understood [16].

The nerve growth factor receptor (NGFR), also known as CD271 or p75 neurotrophin receptor (p75^NTR^) belongs to the tumor necrosis factor receptor (TNFR) superfamily and functions as a low-affinity receptor for all the neurotrophins [17]. Initially, NGFR was studied for its critical role in the embryonic development, where it serves as a marker of neural crest stem cells [17]. Its role in embryonic stem cells has sparked interest in its relevance to cancer, particularly in the context of CSC biology. In various tumor types, including melanoma [18], head and neck squamous carcinomas [19] or breast cancer [20], NGFR^+^ tumor cells possess the ability to initiate and sustain tumor growth *in vivo*. Moreover, NGFR-expressing cells exhibit hallmark CSC traits, such as resistance to chemotherapy [21] or targeted therapies [22]. Recently, attention has shifted to the capacity of NGFR^+^ tumor cells to evade the immune system. Studies in melanoma have demonstrated that NGFR contribute to immune exclusion and ICI resistance through modulation of natural killer (NK) cell activity [23], downregulation of antigens associated with differentiated cells [24] and their colocalization with programmed death-ligand 1 (PD-L1)^+^ tumor cells in regions that evade cytotoxic T cell activity [25].

Here, we evaluated NGFR expression across BLCA models representing both luminal and basal phenotypes. We found NGFR expression in a subpopulation of basal tumor cell lines and organoids exhibiting basal features. Analysis of single-cell RNA-seq data (scRNA-seq) from BLCA patients revealed that NGFR is restricted to basal-like KRT5^+^/KRT14^+^ tumor cells and is absent in luminal UPK3A^+^ cells. Clinically, NGFR serves as an independent poor prognostic marker both in the TCGA and IMvigor210 cohorts both by univariate and multivariate analysis. Importantly, NGFR showed stronger prognostic value than other basal markers. In these cohorts, NGFR expression was associated with stromal cell infiltration, enriched in a subtype of pro-tumoral fibroblasts. In the UROMOL2 cohort, NGFR identifies Class 2b patients, associated with poor prognosis and high stromal content. In the IMvigor210 cohort, NGFR-high tumors showed low response to ICIs, low tumor mutational burden (TMB), reduced neoantigen load, and an immune-excluded phenotype characterized by elevated signature of TGFβ activity in cancer-associated fibroblasts (CAFs) and potential poorer response to immunotherapy. These results were independently validated in the IMvigor010 cohort. Functionally, NGFR knockout (KO) in SCaBER cells reduced stemness-associated programs and invasive capacity, supporting a role for NGFR in aggressive tumor cell behavior. In summary, our study demonstrates that NGFR is a novel marker of a basal-like subpopulation in BLCA with prognostic value highlighting its potential as a therapeutic target.

## Methods

### Immunohistochemistry

Healthy human urothelial tissues were obtained from patients undergoing surgery for non-oncological indications, such as endometriosis, and were provided by the University Hospital Erlangen. All human studies were conducted in accordance with the principles of the Declaration of Helsinki. Informed written consent was obtained from each subject prior to sample collection. The human investigations were performed after approval by the FAU Ethical Committee (approval number 22-343-B). Porcine urothelial samples were collected from the Spanish National Centre for Cardiovascular Research (CNIC, Madrid, Spain). Tissues were fixed in 10% formalin for 24 hours, transferred to ethanol 50% overnight, embedded in paraffin and sectioned at 2.5 µm. Sections were mounted on Superfrost® Plus slides and dried overnight. Stainings were performed using automated platforms (Discovery ULTRA, Ventana-Roche, or Autostainer Link 48, Dako). After antigen retrieval and peroxidase blocking (3% H_2_O_2_), sections were incubated with NGFR antibody (0.2 mg/ml; Abcam; ab3125; RRID:AB_303531) and CK5 (Dako, IR780), followed by HRP-conjugated anti-mouse secondary (ab133469, Abcam; RRID:AB_10695944). For co-localization analyses, adjacent serial sections from healthy human and porcine bladders were stained in parallel with NGFR and CK5. Detection was achieved with 3,3’-diaminobenzidine tetrahydrochloride (DAB) with Envision FLEX, Novolink polymer, or OmniMap Discovery Ultra platforms. Nuclei were counterstained with Harris’ hematoxylin. Positive and negative control tissues were included in each run, and images were acquired by brightfield microscopy. All IHC experiments were replicated in 3 independent samples. Slides were reviewed by experienced pathologists (F.X. Real and E. Caleiras), and IHC analyses were performed in multiple independent human and porcine samples as indicated in the figure legends. For murine tissues (healthy bladder, BBN-induced lesions, organoids, and xenografts), NGFR immunohistochemistry was performed using a rat monoclonal anti-NGFR antibody (NORI146C/E2; Monoclonal Antibodies Core Unit, clone AM (146C)), followed by HRP-conjugated anti-rat secondary antibody and DAB detection, as described above. The animal experimentation was approved by the Autonomous Community of Madrid (PROEX 034.5/21).

### Flow cytometry

All cells were maintained at 37 °C in a humidified atmosphere with 5% CO₂ and subcultured using trypsin-ethylenediaminetetraacetic acid 0.05% (EDTA) (Thermo Fisher Scientific). Cells were detached using a Cell Dissociation Buffer (Gibco™ Cell Dissociation Buffer, enzyme-free, PBS; 11530456), washed twice (PBS, 0.1% BSA, 5mM EDTA), and stained with an NGFR APC-conjugated antibody (Biolegend, ME20.4). NGFR expression was assessed in the following human BLCA cell lines: UM-UC-7, UM-UC-12, RT4, SCaBER, VMCUB1 and TCCSUP. All cells were cultured in Dulbecco’s Modified Eagle Medium (DMEM) mixture supplemented with MEM Non-Essential Amino Acids Solution (#11140-050, Gibco) and 10% FBS. Additionally, gentamicin (Sigma-Aldrich; 200 µg/mL), glutamine (Sigma-Aldrich; 2 mM) and sodium pyruvate (Sigma-Aldrich; 10 mM) were added to all cell lines. BD FACSCanto™ II cytometer was used for acquisition and FlowJo v10.10 (RRID:SCR_008520) for analysis. Singlets were gated using FSC-A and FSC-H, followed by exclusion of dead cells using DAPI (0.5 µg/mL). The percentage of NGFR-positive cells for each cell line was summarized as the mean of three replicate measurements and comparison between basal and luminal cell lines were assessed using a two-sided Welch’s t-test.

### Generation of NGFR-KO cell lines

HEK293T cells maintained in DMEM in 10-cm dishes were co-transfected with 9 µg lentiviral targeting vector (EndoFree Maxi Prep; QIAGEN), 5 µg psPAX2 (Addgene #12260), 3 µg pMD2.G (Addgene #12259), and 62.5 µl Lipofectamine 2000 (Life Technologies). Viral supernatants were collected at 48 h and filtered (0.45 µm low protein-binding; Millipore). Target cells were transduced in suspension at low MOI with viral supernatant diluted 1:2 in the presence of 8 µg/ml polybrene, then plated at 1×10^5 cells per well in 6-well plates. SCaBER NGFR knockout lines were generated using CRISPR/Cas9. sgRNAs targeting human NGFR (sgNGFR1 sequence: 5’-GGTAGTAGCCGTAGGCGCAG-3’ and sgNGFR2 sequence: 5’-GTGTGGACCGTGTAATCCAA-3’) were cloned into lentiCRISPRv2 (Addgene #52961). Control sgRNAs targeted human AAVS1 (5′-CCTCTAAGGTTTGCTTACGA-3′). Lentiviruses were generated and cells transduced as above, followed by puromycin selection (1 µg/ml; InvivoGen). Knockout populations were enriched by sorting for loss of NGFR expression (BD Influx) and validated by flow cytometry (FACS Canto) using an NGFR APC-conjugated antibody (Biolegend, ME20.4)

### Generation of tumor spheroids

The spheroids were generated by culturing SCaBER cells under suspension growth conditions. The cells were seeded in ultra-low attachment 6-well plates at a density of 3,000 cells per well for each experimental condition (SCaBER control, KO1, and KO2). Cells were cultured with serum-free DMEM/F12 supplemented with B27 (1:50), EGF (10 ng/mL), and bFGF (10 ng/mL) and were maintained under these conditions for 2 weeks to promote spheroid formation. After 2 weeks, spheroids were captured with the Leica THUNDER microscope.

For the analysis, quantification was performed using a custom Python pipeline from Leica .lif files (merged images). Images were converted to grayscale (normalized to [0–1]) and smoothed with a Gaussian filter (σ=2). Spheroids were segmented using Otsu thresholding (inverted when required), followed by morphological closing (disk radius 3), removal of small objects (<200 px), and exclusion of objects touching image borders. Objects were retained as spheroids only if they met strict morphological criteria (area 800–20,000 px, solidity >0.9, eccentricity <0.8). For each spheroid, area, equivalent diameter, and mean intensity were extracted and exported for downstream analysis.

### Transwell migration assay

SCaBER control (CTL) and NGFR knockout clones (KO1, KO2) were assessed in Transwell inserts under serum-starved conditions to promote migration. Cells were maintained in DMEM supplemented with MEM non-essential amino acids and 10% FBS, and migration was allowed to proceed for 24 h. Membranes were stained with DAPI (5 µg/mL, 20 min) and imaged by confocal microscopy (Leica SP5 MP). For each condition, duplicate wells were analyzed; four fields per well were acquired as z-stacks (62 planes).

The analysis was performed using the confocal z-stacks and a custom Python pipeline. For each field, DAPI (C00) was segmented in 3D after Gaussian smoothing and Otsu thresholding. Objects were hole-filled, small objects were removed, and nuclei were separated by distance-transform watershed seeded. For each nucleus, the z-centroid was computed. The membrane position was estimated from the second channel (C01) as the peak in mean intensity along z. Cells with z-centroids beyond this peak were considered migrated. Fields with <800 total nuclei were excluded. Counts were averaged across fields, then across duplicate wells to obtain condition-level values.

### BBN-induced murine bladder tumors, organoids, and xenografts

Bladder tumors were generated in C57BL/6 mice by exposure to the carcinogen N-butyl-N-(4-hydroxybutyl) nitrosamine (BBN) supplied in the drinking water at a concentration of 0.025%. Drinking water was replaced twice weekly, and treatment was maintained for up to 20 weeks. With this regimen, mice developed carcinoma in situ at approximately 10–12 weeks and invasive carcinomas at 20–25 weeks. Bladders bearing BBN-induced tumors were collected at the malignant stage and processed to establish three-dimensional tumor organoids, following protocols previously described for mouse bladder organoids [65]. Organoids derived from BBN-induced tumors were subsequently dissociated into two-dimensional (2D) cultures. These BBN 2D cells were expanded *in vitro* and injected subcutaneously into the flank of Nude/Nude immunodeficient mice to generate xenograft tumors. Xenografts were harvested at endpoint, fixed in formalin, embedded in paraffin, and processed for NGFR immunohistochemistry as described above.

### CCLE bladder cancer cell line transcriptomic analysis

Transcriptomic data and annotations for BLCA cell lines were obtained from the CCLE [26]. We first selected all urothelial carcinoma cell lines (n = 37) and classified them into molecular subtypes according to the consensus MIBC classification [6], resulting in 18 Basal/Squamous (Ba/Sq), 9 Luminal Papillary (LumP), 2 Luminal Unstable (LumU), and 8 Neuroendocrine-like (NE-like) lines. For NGFR expression analyses comparing luminal versus basal phenotypes, we included the 18 Ba/Sq and 11 luminal (LumP + LumU) lines and excluded NE-like models.

To characterize the NGFR⁺ basal phenotype, we restricted downstream analyses to the 18 Ba/Sq BLCA cell lines. Gene-level expression values were used to compute Pearson correlation coefficients between NGFR and all other genes in the genome, generating a ranked list from the most positively to the most negatively correlated genes. This ranked list was then used as input for a pre-ranked gene set enrichment analysis (GSEA) using Reactome pathways. Pathways were considered significantly associated with NGFR expression based on normalized enrichment score (NES) and false discovery rate (FDR)–adjusted q-values. For correlation analyses with specific genes, we grouped markers into three categories: basal-like markers, epithelial–mesenchymal transition (EMT) markers, and invasion-related markers. Basal markers were selected based on established BLCA molecular subtype studies [6, 27], EMT markers were derived from canonical EMT reviews [28], and invasion markers were taken from published invasion/metastasis gene sets in urothelial carcinoma [29].

### Patient derived organoids

The analysis of NGFR expression in patient-derived organoids was conducted using a repository of published expression data [30], comprising 16 human organoids derived from bladder tumors (both NMIBC and MIBC). NGFR expression was assessed in organoids categorized in the publication as either luminal or basal, excluding those classified as mixed.

### Patient cohort scRNA-seq

For the scRNA-seq analysis of patient samples, we used published data [31], which includes eight primary bladder urothelial tumors (two low-grade and six high-grade) along with three adjacent normal mucosa samples. Data preprocessing and unsupervised clustering were carried out to distinguish cell types and individual patients. Single cell data was analyzed from FastQ files and pseudoaligned to the GRCh38 cDNA sequence using kallisto bustools. Individual bus files were converted into count matrices using the BUSpaRse package and analyzing using Seurat (RRID:SCR_016341) and Scanpy (RRID:SCR_018139) toolkits. Clusters were grouped as epithelial or TME cells, excluding non-tumor mucosa. NGFR and other markers (KRT5, KRT14, UPK3A) were visualized by Uniform Manifold Approximation and Projection (UMAP).

### Patient cohorts bulk RNA-seq

The relevance of NGFR was analyzed in well-annotated large patient cohorts. We used TCGA [5] and IMvigor210 [32], UROMOL2 [4] and IMvigor010 [33]. The TCGA cohort includes 411 MIBC patients, 407 with expression data utilized to examine NGFR expression in relation to overall survival, tumor subtypes and the TME. TCGA data (BLCA cohort) was downloaded from the Genomic Data Commons (GDC) portal (https://portal.gdc.cancer.gov/). IMvigor210 contains 348 MIBC patients, all with survival and expression data. UROMOL2 included 535 NMIBC patients with expression profiles. IMvigor010 includes 728 MIBC patients with expression profiles [33]. IMvigor210 and UROMOL2 datasets were accessed via [4, 32].

### Survival analysis

Univariate survival analysis was conducted using Kaplan-Meier analysis and log-rank test by GraphPad Prism (RRID:SCR_002798). In both the TCGA and IMvigor210 cohorts, patients were classified into quartiles based on NGFR expression levels: Q1 (lowest quartile of expression), Q2 (low-mid), Q3 (mid-high), and Q4 (very high). Clinical covariates for multivariate Cox models were selected based on clinical relevance and data availability in each cohort. Multivariate cox models included clinical covariates: TCGA included age, sex, disease stage, histologic grade and radiation history; IMvigor210 included sex, baseline ECOG score, metastasis status, BCG, platinum therapy, tobacco history (ever vs. never), tissue origin (non-bladder vs. bladder) and smoking. Multivariate with other basal markers included KRT5, KRT14 and CD44. Significance was evaluated using the Wald test. Survival was evaluated using Python package (RRID:SCR_016074). For Cox proportional hazards multivariate analysis, results are presented as hazard ratios (HR) with 95% confidence intervals, and the proportional hazards assumption was assessed using Schoenfeld residuals. In both the TCGA-BLCA and IMvigor210 cohorts, overall survival (OS) was used for survival analyses. In the IMvigor010 cohort, disease-free survival (DFS) was defined as the time to the first occurrence of local (pelvic) or urinary tract recurrence, distant urothelial carcinoma metastasis, or death from any cause. Patient age was not available in the downloaded IMvigor210 clinical table; therefore, it could not be included in the multivariable model. We also reviewed neoadjuvant therapy information in TCGA-BLCA, but this variable was present in only a small number of cases and was therefore not included in the multivariable model. Age and NGFR expression were modeled continuously. Sex, radiation therapy, intravesical BCG administration, platinum treatment, tobacco history, and tissue origin were modeled as binary variables. Baseline ECOG score, AJCC stage, histologic grade, and metastatic disease status were modeled according to the coding used in the Cox models and are specified in the figure labels.

### Consensus MIBC subtype analysis

MIBC subtype described by Kamoun et al. [6] were assign to each patient in TCGA and IMvigor210 using an R package (https://github.com/cit-bioinfo/consensusMIBC). For quartile analysis patients were grouped by NGFR expression as previously described.

### Ecosystem deconvolution

We applied the Ecotyper Carcinoma tool (https://ecotyper.stanford.edu/) to evaluate TME composition, identifying ten conserved ecosystem types and associated immune and stromal cell subtypes.

### Immune phenotype *in silico* classification

Immune phenotypes (inflamed, excluded, or desert) were obtained from the IMvigor210 dataset as defined in the original publication, which classifies tumors based on the spatial distribution of tumor-infiltrating lymphocytes [32]. In parallel, Ecotyper was applied to infer tumor ecosystems from bulk RNA-seq data [34]. To further corroborate the immune-excluded phenotype, transcriptomic markers of fibroblast activation and TGFβ signaling were evaluated in NGFR-high tumors [32].

### ICIs resistance conditionings

Resistance mechanisms were explored in IMvigor210 [32], using TMB (n=272) and neoantigen levels (n=245). For CD8^+^ T effector cell and TGFβ signaling signatures in CAFs, we applied the gene signatures previously described in the referenced study [32]. Scores for each patient in both signatures were calculated based on transcriptomic profiles using single-sample gene set enrichment analysis (ssGSEA) with a gene set variation analysis package (GSVA; https://github.com/rcastelo/GSVA; RRID:SCR_021058)

### Data availability

Code is publicly uploaded and available (https://github.com/jgagullo). The data will be made available from the corresponding author upon reasonable request.

## Results

### NGFR expression is associated with the basal phenotype in BLCA

To assess NGFR expression in the normal urothelium, we examined its localization by immunohistochemistry. We found that NGFR expression in urothelium is confined to the basal layer in human bladder and co-localizes with the basal keratin KRT5 (**Fig. 1A, Supplementary Fig. 1A–D**). We observed the same pattern in porcine bladder, where NGFR⁺ cells overlap with KRT5 expression in serial sections (**Supplementary Fig. 1E**), in line with previous reports [35].

**Figure 1.**
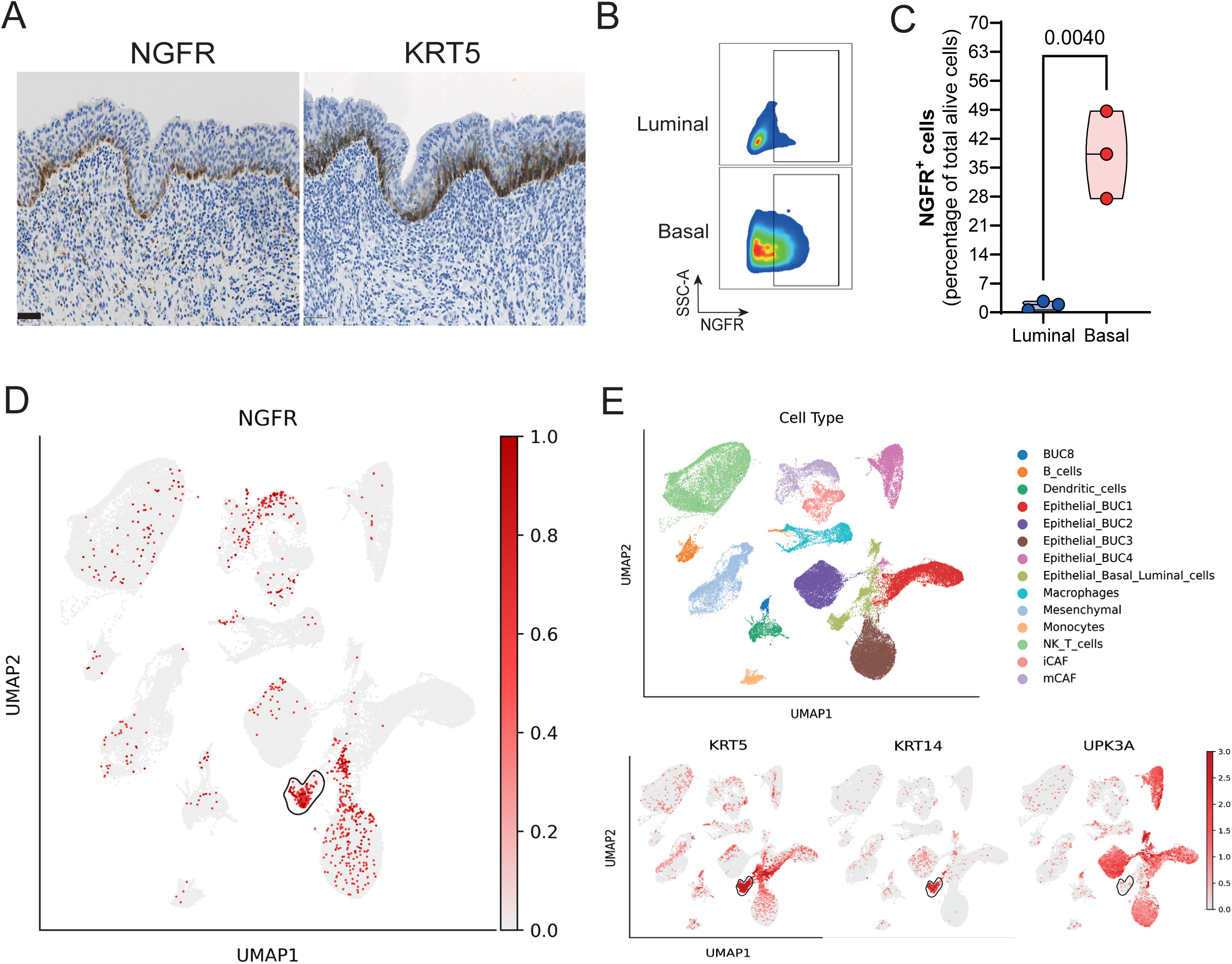
NGFR identifies a basal cell subcluster in human BLCA. (**A**) Co-localization of NGFR with basal keratins in healthy human bladder tissue. Representative serial sections of the same healthy urothelium stained for NGFR and KRT5 are shown. Scale bar: 50 µm. Sample size: n = 4 healthy bladders stained for NGFR expression and n = 2 healthy bladders stained for KRT5 (see Supplementary Fig. S1A–D for NGFR staining in all samples). (**B, C**) Flow cytometry analysis of NGFR expression in a panel of human bladder cancer cell lines. The panel includes three luminal lines (UM-UC-7, UM-UC-12, and RT4) and three basal lines (SCaBER, VMCUB1 and TCCSUP). Each measurement was performed in three independent experiments. Representative density plots from one experiment are shown in **B**, while aggregated quantifications are also presented in **C**. Each point in **C** represents one cell line, plotted as the average of the three independent experiments, and boxplots summarize the distribution in each cell subtype group. P values were calculated using a two-sided unpaired Welch’s t-test comparing the two cell subtype groups (luminal and basal). (**D, E**) UMAPs of the scRNA-seq dataset from BLCA patients [31]. NGFR expression in each cluster is shown (**D**), as well as cell type assignment, differentiating tumor cells from immune cells, and expression of luminal (UPK3A) and basal (KRT5 and KRT14) markers (**E**). Within the cluster annotated in the original dataset as Epithelial_Basal_luminal_cells, the black-circled subpopulation corresponds to basal-like epithelial cells, showing enrichment for NGFR, KRT5 and KRT14, together with low UPK3A expression.

We next assessed NGFR expression in human bladder tumor cells, where its role remains uncharacterized. To do so, we analyzed NGFR levels in a panel of human cell lines by flow cytometry. The panel included three luminal cell lines (UM-UC-7, UM-UC-12, and RT4) and three basal lines (SCaBER, VMCUB1 and TCCSUP) [36]. NGFR expression was upregulated in basal cell lines compared with luminal models (**Fig. 1B, C**). To confirm these findings in a larger set of cancer cell lines, we interrogated the Cancer Cell Line Encyclopedia (CCLE) [26]. NGFR expression was mainly observed in basal cell lines: while not all basal cells expressed NGFR, we found that all cell lines with high NGFR levels mostly belonged to the basal subtype (**Supplementary Fig. 1F**). This pattern of expression was further confirmed by the analysis of published human organoid data [30], which showed elevated NGFR levels in basal organoids compared to luminal types (**Supplementary Fig. 1G**).

To further characterize the NGFR⁺ basal phenotype in BLCA tumors cells, we analyzed bulk RNA-seq data available in the CCLE database [26]. Analysis using GSEA, based on correlation with NGFR expression, revealed that the main upregulated pathways were related to epithelial differentiation and keratinization (**Supplementary Fig. 1H, I**). When evaluating the correlation between NGFR and different groups of CSC-like markers (including basal-like phenotype, EMT, and invasion markers) we observed that the strongest positive association was with basal markers (**Supplementary Figure 1J**), including KRT14, a canonical basal keratin. In contrast, no correlation was detected with classic EMT or invasion markers (**Supplementary Figure 1J**). Taken together, these data support the notion that NGFR defines a basal epithelial state within Ba/Sq cells, independent of mesenchymal or invasion programs. To confirm if NGFR expression is restricted to basal BLCA cancer cells in a clinical setting, we evaluated a scRNAseq dataset of tumors derived from BLCA patients [31]. Our analysis revealed that NGFR expression is predominantly confined to tumor cells expressing KRT5 and KRT14, while being absent in the luminal UPK3A^+^ regions (**Fig. 1D, E**), confirming the results that we previously obtained in cellular models. NGFR expression was also detected more diffusely in other epithelial and stromal cell clusters.

Given that our initial analyses showed that NGFR expression is associated with the basal phenotype, we evaluated its functional contribution to stemness and invasion. We generated NGFR knockout (NGFR-KO) cells in the basal SCaBER line—chosen for its high endogenous NGFR expression—using two independent CRISPR guide RNAs. NGFR depletion was validated by flow cytometry (**Supplementary Fig. 2A**). We found that NGFR-KO reduced spontaneous primary spheroid formation under suspension conditions (**Supplementary Fig. 2B, C**), demonstrating that NGFR contributes to spheroid growth capacity in this model. In addition, we evaluated the migratory capacity of these cells by transwell assay. We observed that NGFR-KO reduced the migratory capacity of the tumor cells (**Supplementary Fig. 2D, E**). To test the relevance of NGFR in *in vivo* models, we analyzed its expression in a murine model treated with the carcinogen BBN, which specifically generate basal/squamous malignant lesions in the urinary bladder [7, 8, 37]. In this context, NGFR showed minimal expression in healthy C57BL/6 mouse urothelium, increased in BBN-induced non-malignant lesions, and increased further in malignant-stage lesions (**Supplementary Fig. 2F**). Furthermore, NGFR expression was maintained in organoids that were generated from the BBN-derived malignant lesions (**Supplementary Fig. 2G**) and in secondary xenograft tumors obtained after subcutaneous inoculation of BBN tumorigenic cells (**Supplementary Fig. 2H**). Taken together, these results suggest that NGFR is induced during tumor progression in murine basal BLCA, suggesting that NGFR plays a functional role in characteristics associated with the stem-like and invasive phenotype in BLCA.

### NGFR expression is an independent prognostic marker in BLCA

Our results indicate that NGFR serves as a marker for a subset of cells within the basal population. The basal phenotype in BLCA is generally associated with poor prognosis compared to luminal subtypes [5]. Kaplan-Meier survival analyses were performed using quartiles of NGFR expression: Q1 representing the lowest levels of NGFR expression and Q4 representing the highest expression, with Q2 and Q3 as intermediate values. For multivariate analysis, we used continuous NGFR expression. To evaluate the relevance of NGFR in BLCA survival we first interrogated The Cancer Genome Atlas (TCGA) cohort, which is a heterogeneous patient dataset representing the diversity of the disease [5]. We found that stratifying TCGA BLCA patients by NGFR expression quartiles is associated with poor overall survival (OS), with Q1 having the best survival relative to the rest (**Fig. 2A**). However, the survival pattern across Q2–Q4 was not strictly ordinal, and the most evident difference was observed between Q1 and the remaining quartiles. Consistent with this observation, an analysis comparing Q1 versus Q2–Q4 showed that patients in the lowest quartile of NGFR expression had significantly better survival than the remaining patients (**Fig. 2B**). Importantly, we observed using Cox proportional hazards that the prognostic effect of NGFR was independent of other relevant clinical factors included in the TCGA dataset such as age, sex, tumor stage, histological grade and previous radiation treatment (**Fig. 2C**). In this multivariable model, NGFR was associated with OS as a continuous variable (HR 1.22, 95% CI 1.05–1.42, p-value = 0.012), and no evidence of violation of the proportional hazards assumption was observed for NGFR (Schoenfeld test p = 0.536). To contextualize NGFR within the basal program, we compared its prognostic association with that of canonical basal markers using the same multivariable clinical framework. In TCGA, KRT5 and KRT6B also showed significant associations in these gene-by-gene multivariable models, whereas KRT14 and CD44 did not (**Supplementary Fig. 3A**). When comparing all the basal markers in a multivariate model together, NGFR was the only one that maintained the prognostic effect (**Supplementary Fig. 3B**). Additionally, Spearman correlation analysis showed that NGFR was only weakly correlated with these canonical basal markers, whereas the classical basal markers were more strongly correlated with each other (**Supplementary Fig. 3C**).

**Figure 2.**
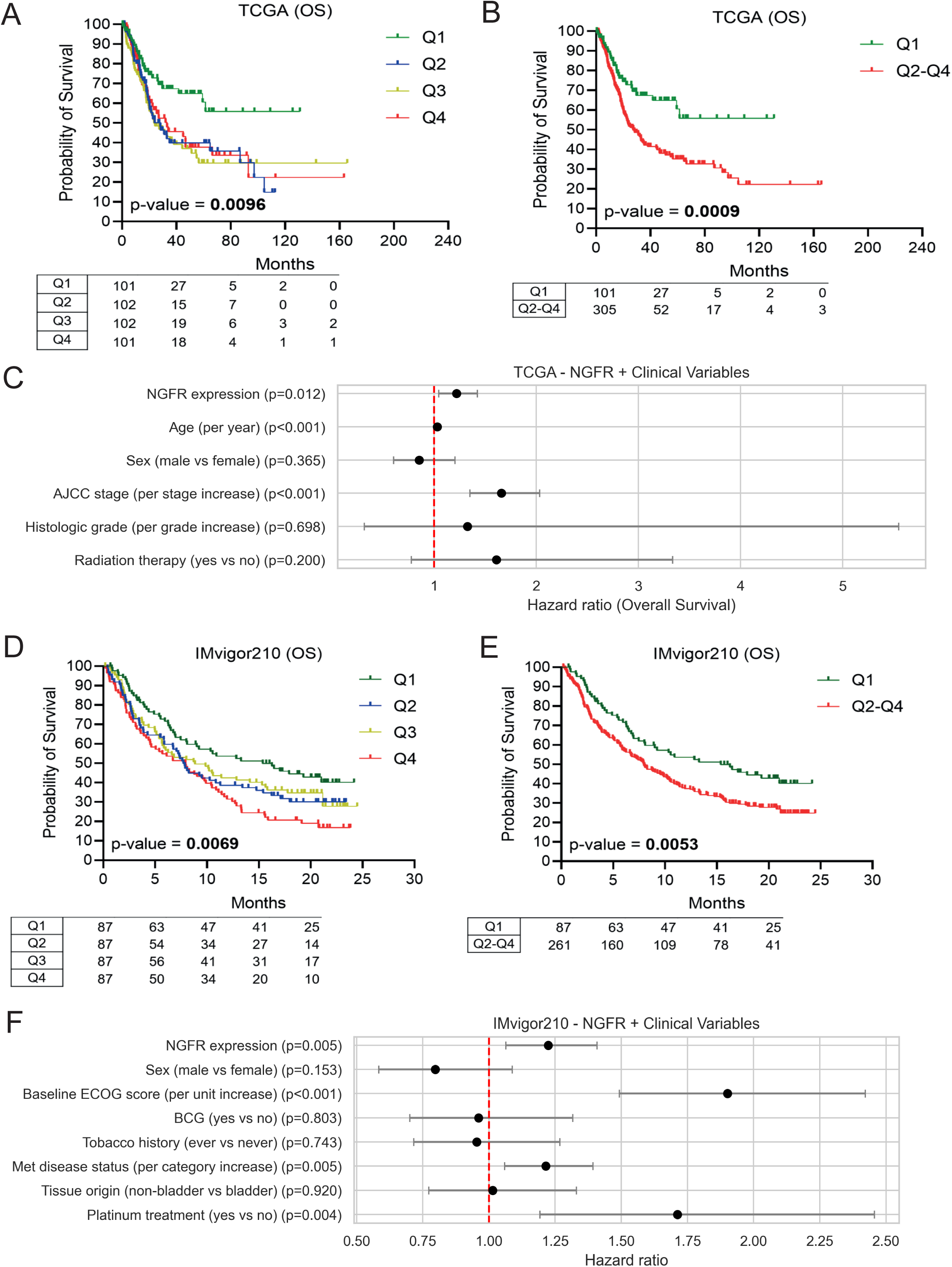
NGFR expression is a poor prognostic marker in BLCA patients. (**A–C**) Survival analysis in the TCGA BLCA cohort. Univariate analysis by Kaplan-Meier curves according to NGFR expression quartiles (**A**), dichotomized Kaplan-Meier analysis comparing Q1 versus Q2–Q4 (**B**), and multivariate analysis by Cox proportional hazards model (**C**) are shown. (**D–F**) Survival analysis in the IMvigor210 cohort [32]. Univariate analysis by Kaplan-Meier curves according to NGFR expression quartiles (**D**), dichotomized Kaplan-Meier analysis comparing Q1 versus Q2–Q4 (**E**), and multivariate analysis by Cox proportional hazards model (**F**) are shown. For Kaplan-Meier analysis, patients were grouped according to quartiles of NGFR expression, with Q1 having the lowest expression and Q4 the highest. In the TCGA cohort, Q1 included 101 tumors and Q2–Q4 included 306 tumors in the dichotomized analysis; in the IMvigor210 cohort, each quartile included 87 tumors, and Q2–Q4 included 261 tumors in the dichotomized analysis. The number of patients at risk is shown below each Kaplan–Meier curve. For the Cox models, NGFR expression was evaluated as a continuous variable and results are shown as hazard ratios (HR) with 95% confidence intervals, adjusting for the main clinical characteristics of each cohort. In IMvigor210, the multivariable model included baseline ECOG score in addition to the other clinical covariates. Tobacco history and tissue origin were modeled as ever vs. never smoking and non-bladder vs. bladder tissue origin, respectively. P-values were calculated using the log-rank test for the Kaplan-Meier analysis, and by Wald test for the Cox model.

Additionally, we applied the same approach to a second BLCA cohort. For this purpose, we selected IMvigor210 [32], a cohort enriched in metastatic patients treated with ICIs. Importantly, we found that stratifying patients based on quartiles of NGFR expression was sufficient to find differences in overall survival (OS): Q1 patients had the best OS, while Q4 patients had the worst, with Q2 and Q3 representing intermediate prognostic state (**Fig. 2D**). Consistently with the TCGA cohort, a dichotomized analysis comparing Q1 versus Q2–Q4 showed that patients in the lowest quartile of NGFR expression had significantly better survival than the remaining patients (**Fig. 2E**). Importantly, we confirmed that NGFR effect on survival is also independent of other relevant clinical characteristics collected in the cohort such as sex, baseline ECOG score, BCG treatment, tobacco smoking, metastatic status, tissue analyzed, or platinum treatment in IMvigor210 (**Fig. 2F**). In this multivariable model, NGFR was associated with OS as a continuous variable (HR 1.21, 95% CI 1.06–1.41, p-value = 0.005), and no evidence of violation of the proportional hazards assumption was observed for NGFR (Schoenfeld test p = 0.130). Applying the same approach in IMvigor210 to that applied in the TCGA cohort, we evaluated the correlation of canonical basal markers with survival in a multivariate model.

NGFR, KRT5 and KRT6B were significantly associated with poor survival, whereas KRT14 and CD44 were not (**Supplementary Fig. 3D**). As in TCGA, NGFR was the only marker that maintained the effect in poor prognosis in a multivariate model including all markers (**Supplementary Fig. 3E**). Additionally, correlation analysis showed that NGFR was only weakly correlated with the canonical basal markers, while stronger correlations were observed among the classical basal markers themselves (**Supplementary Fig. 3F**). These results suggest a role for NGFR in bladder cancer, being associated with poor prognosis independently of clinical characteristics and other basal markers in two independent cohorts analyzed.

### NGFR expression defines stromal-enriched and basal-like tumor subtypes in BLCA

To further characterize the potential role of NGFR in BLCA, we analyzed its expression across different subtypes. Specifically, we analyzed the association between NGFR expression quartiles and consensus tumor MIBC subtypes [6], within two independent cohorts: TCGA and IMvigor210 (**Fig. 3A, B**). In both cohorts, our analysis revealed a shift as NGFR expression increased. Notably, high NGFR expression (found in the upper quartiles, Q3 and Q4) was more common among the basal-like and stroma-rich subtypes, typically associated with aggressive MIBC [6] and was less prevalent among the luminal subtypes, particularly luminal papillary (LumP) and luminal unstable (LumU). Similarly, we evaluated the associations of NGFR and NMIBC. To address specifically NMIBC, we interrogated the latest UROMOL2 dataset [4]. Notably, we observed high NGFR expression (Q3 and Q4) was enriched among NMIBC Class 2b tumors (**Supplementary Fig. 4A**). Class 2b is defined as poor prognostic subtype, with stem-like features and a high stromal compartment [3, 4]. We also evaluated the immune-score of these tumors and observed that NGFR expression was positively associated with high immune infiltration (**Supplementary Fig. 4B**). These results show that NGFR, both in MIBC and NMIBC, identifies tumors with a high stromal compartment.

**Figure 3.**
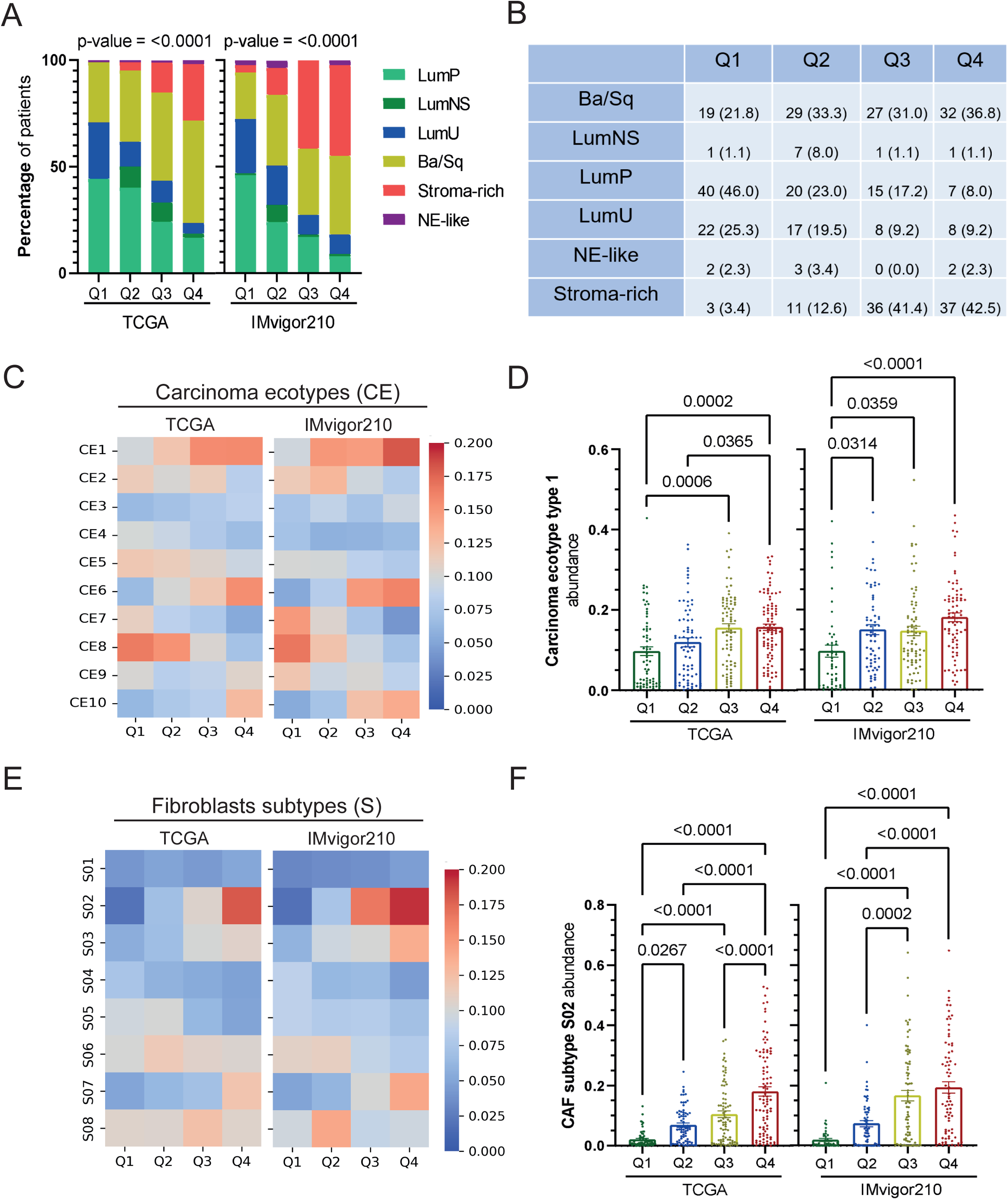
BLCAs with high NGFR expression are associated with an increased stromal compartment. (**A**) Distribution of consensus molecular subtypes [6] as a function of NGFR expression quartiles in the TCGA cohort (left) and in the IMvigor210 cohort (right). P-values were calculated by Chi-square test (and Fisher’s exact test), comparing the number of patients in each group (the corresponding percentage is represented). (**B**) Summary table with the number and percentage (in parentheses) of patients in each molecular subtype in the IMvigor210 cohort. (**C, D**) Analysis by Ecotyper [34] in TCGA (left) and IMvigor210 (right) patients, according to quartiles of NGFR expression. Heatmaps of the 10 tumor ecosystems defined by Ecotyper (**C**), and barplots of the CE1 ecosystem (**D**) are shown. (**E, F**) Deconvolution of fibroblasts by Ecotyper according to quartiles of NGFR expression in both TCGA (left) and IMvigor210 (right) cohorts. Heatmaps of all fibroblast subtypes (**E**), and barplots of the CAF S02 subtype (**F**) are presented. P-values for barplots were calculated by ordinary one-way ANOVA.

To gain deeper insights into the role of NGFR in the TME, we conducted an ecosystem analysis using the Ecotyper tool in both TCGA and IMvigor210 cohorts [34]. This tool categorizes carcinoma ecotypes (CE) based on conserved immune and stromal profiles. Our analysis revealed that high NGFR expression is associated with carcinoma CE1 and CE6 in both cohorts (**Fig. 3C**). Specifically, CE1 was the most frequent in NGFR Q4 patients (**Fig. 3D**). Both CE1 and CE6 are characterized by a significant stromal component enriched in fibroblasts and a reduced presence of immune cells [34]. Further analysis of the fibroblast subtypes within these ecosystems revealed a significant enrichment of the S02 subtype in tumors with high NGFR expression in both cohorts (**Fig. 3E, F**). The S02 fibroblast subtype is formed by CAFs expressing extracellular matrix (ECM) regulatory proteins and TGF-β signaling molecules, among other features [38].

Given this association between NGFR and a fibroblast-rich microenvironment, we next explored the relationship between NGFR and its canonical ligands at different levels of biological resolution. First, we analyzed basal bladder cancer cell lines from the CCLE dataset [26]. Among the neurotrophin ligands, only NTF4 displayed a correlation with NGFR (**Supplementary Fig. 5A**). Analysis in bulk RNA-seq data from primary MIBC tumors (TCGA-BLCA) [5] showed a moderate positive correlation between the expression of NGFR and NGF, BDNF and NTF3 (**Supplementary Fig. 5B**). This pattern was reproduced in the IMvigor210 cohort [32] of advanced MIBC patients treated with ICIs (**Supplementary Fig. 5C**). Since these analyses include both tumor and stromal cells, we analyzed scRNA-seq data from bladder cancer in order to define the potential source [31]. These analyses revealed that NGF, BDNF and NTF3 ligands are mainly expressed in CAFs and are virtually absent in tumor epithelial cells (**Supplementary Fig. 5D, E**). Together, these results suggest a CAF-mediated paracrine signaling which aligns with the association between NGFR expression and the stromal-enriched carcinoma ecotypes found in our analysis.

### NGFR expression is associated with poor response to ICIs and an immune-excluded tumor phenotype in BLCA

The role of fibroblasts in MIBC has been studied, particularly in relation to their interactions with the immune system and their contribution to resistance to immunotherapy [32, 39]. Given that NGFR expression is enriched in tumor ecosystems associated with immune cell exclusion and stromal cell enrichment, we investigated its potential as a marker of ICIs resistance. To this end, we analyzed NGFR expression and its correlation with ICIs response in IMvigor210. We found that stratifying patients by NGFR expression was associated with different responses to ICIs (**Fig. 4A**). Specifically, patients in the higher NGFR quartile (Q4) exhibited the lowest response rates to ICIs, suggesting that NGFR expression may contribute to the reduced efficacy of IT. These findings highlight the potential of NGFR as a biomarker associated with poor response to ICIs.

**Figure 4.**
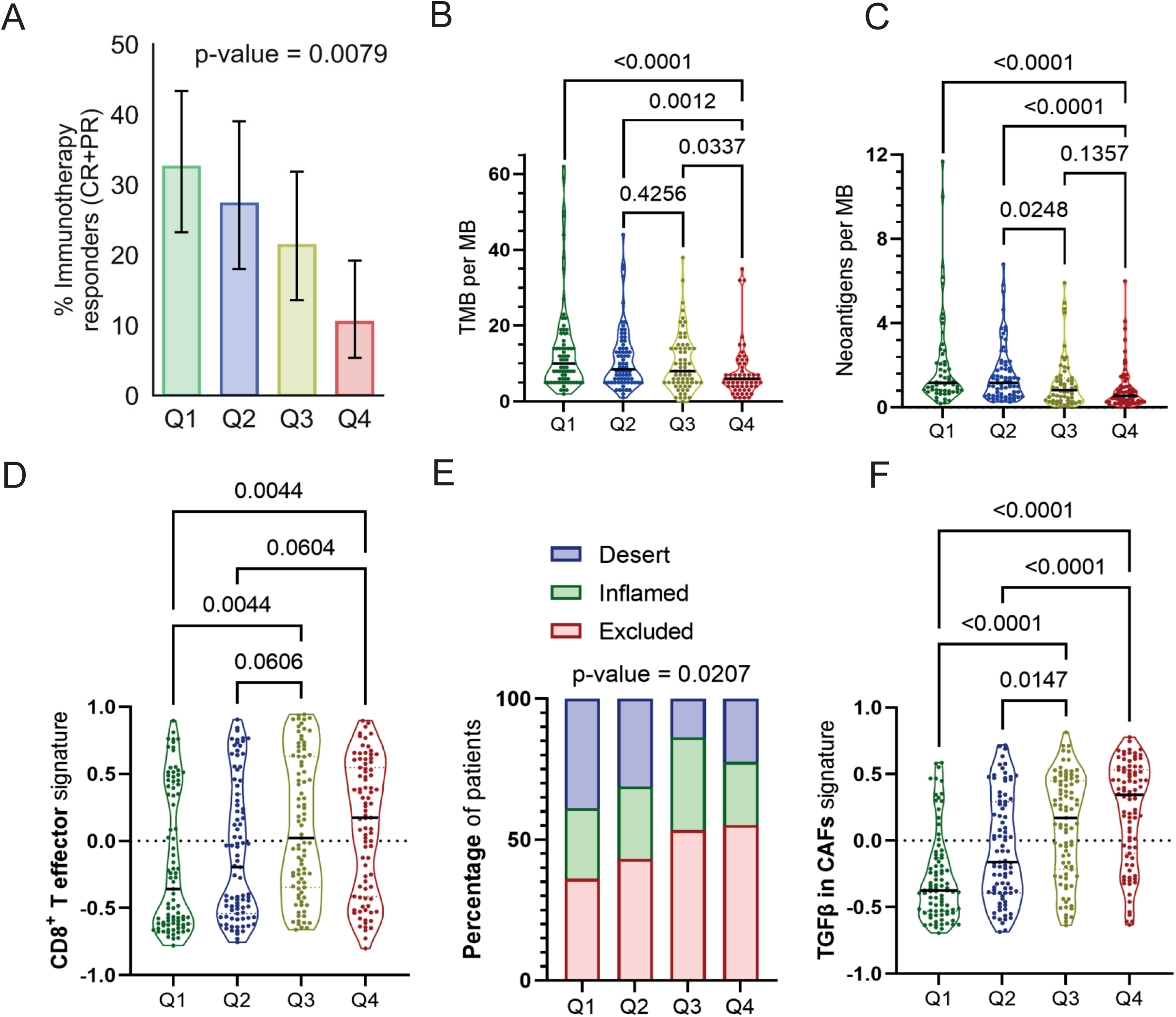
NGFR is a marker of immunotherapy resistance in IMvigor210. (**A**) Percentage of responders to immunotherapy as a function of NGFR expression quartiles in IMvigor210 cohort. P Error bars indicate 95% binomial confidence intervals for the proportion of responders in each quartile. The p value corresponds to the overall comparison of response proportions across the four quartiles using the Chi-square test. (**B, C**) Association between NGFR quartiles and TMB (**B**) and the number of neoantigens (**C**). (**D, E**) Assessment of CD8^+^ T cells in BLCA tumors by effector score (**D**) and immune subtype classification (**E**). (**F**) Association of NGFR expression with TGF-β signaling signature in CAFs. For panels **B** and **C**, p values correspond to two-sided Wilcoxon rank-sum tests for the pairwise comparisons indicated in the plots, with Holm correction for multiple comparisons. P values for gene signatures were calculated by ordinary one-way ANOVA.

To further characterize the NGFR-associated tumor phenotype, we evaluated in IMvigor210 several key factors that are known to influence ICIs resistance in BLCA [32]. This analysis aimed to provide deeper insight into how NGFR expression affect the immune landscape and contribute to resistance mechanisms in these tumors. We observed lower TMB values in tumors from the highest NGFR quartile (Q4) compared with the rest of the quartiles (**Fig. 4B**). Similarly, tumors in the Q4 quartile showed lower neoantigen load than those in the lowest quartile (Q1 & Q2) (**Fig. 4C**). Despite the lower TMB and neoantigen load, tumors in the highest NGFR quartile (Q4) showed higher CD8+ effector signature values than those in the lowest quartile (Q1) (**Fig. 4D**). However, this increase did not translate into effective immune infiltration since NGFR expression identified a higher percentage of patients with an “excluded” immune phenotype, in which CD8^+^ T cells are confined to the tumor periphery, unable to infiltrate the tumor core (**Fig. 4E**). Furthermore, high NGFR expression correlated with a TGFβ signature in activated CAFs (**Fig. 4F**) that has been previously linked to immune exclusion, preventing CD8^+^ T cell infiltration [32]. To further assess these relationships without stratifying NGFR expression into quartiles, we performed correlation analyses using NGFR as a continuous variable. In agreement with the quartile-based analyses, NGFR expression showed a significant inverse correlation with TMB and neoantigen burden, and a positive correlation with CD8+ effector signature and TGFβ signature in CAFs (**Supplementary Fig. 5F-I**). Very similar results were observed in UROMOL2, in which higher NGFR expression quartiles positively correlated with a CD8⁺ T cell signature (**Supplementary Fig. 4C**) and TGF-β signature in CAFs, (**Supplementary Fig. 4D**), showing that these findings are also transferable to NMIBC.

To validate these findings of immune exclusion in an independent cohort, we evaluated the IMvigor010 study, which includes MIBC patients treated with ICIs and has transcriptomic data [33, 40]. In the analysis of the entire cohort, we observed that patients with high NGFR expression had significantly shorter disease-free survival (DFS), supporting the adverse prognostic value of NGFR (**Supplementary Fig. 6A**). Focusing exclusively on the ICI-treated arm, a trend toward shorter DFS and a higher proportion of patients with recurrence was observed in the higher NGFR quartiles, although these differences were not statistically significant (**Supplementary Fig. 6B, C**). Analysis of molecular subtype distribution revealed that elevated NGFR levels were associated with progressive enrichment of the stroma-rich subtype and a reduction in luminal tumors (**Supplementary Fig. 6D**), replicating the pattern observed in TCGA and IMvigor210. Consistently, NGFR showed an inverse association with TMB (**Supplementary Fig. 6E**) and a positive and gradual correlation with the TGFβ signature in CAFs, which increased stepwise across NGFR expression quartiles (**Supplementary Fig. 6F**). Together, these results support in an independent cohort that NGFR is associated with an immunosuppressive stromal tumor phenotype, characterized by low mutational burden and high TGFβ signaling and further support its adverse prognostic value, suggesting a potential association with poorer response to immunotherapy.

## Discussion

Tumor heterogeneity in BLCA presents a critical challenge, particularly regarding resistance to ICIs. Basal cells have gained prominence as a subpopulation with CSC traits, involved in tumor growth and therapy resistance [41]. Under physiological conditions, basal cells regenerate the urothelium upon injury. Recently, immune regulation has been recognized as one of the hallmarks of cancer stemness [16, 42]. CSCs have unique capabilities to evade from immune cells, making them crucial targets for therapeutic approaches [16, 42]. However, the mechanisms behind CSC-mediated immune evasion and ICIs resistance are still not well understood.

We demonstrate that NGFR marks a subpopulation of basal cells in BLCA human models and patient datasets. Whether NGFR expression is a consequence of a basal program or plays a driver role remains an open question requiring further investigation. Evidence from other tumors strongly suggests a driver role for NGFR. In different tumor types such as melanoma, HNSCC and breast cancer, NGFR^+^ cells have been shown to preferentially generate tumors compared to NGFR^-^ cells [18–20]. Furthermore, NGFR^+^ cells are increasingly associated with hallmark features of CSCs, including EMT and invasiveness [43, 44], therapy resistance [21, 22], poor prognosis, and enhanced metastatic capacity [45]. In line with these observations, our *in vitro* loss-of-function experiments indicate that NGFR contributes functionally to a stem-like and invasive phenotype in BLCA, as evidenced by reduced spheroid formation and migration in NGFR-KO cell lines. Additionally, we observed that, beyond being a marker of basal cells within tumors, NGFR expression holds prognostic significance in patients independently of clinical features and other basal markers. In survival analyses, the most consistent difference across cohorts was observed between tumors in the lowest quartile of NGFR expression and the remaining cases, rather than as a strictly proportional gradient across all quartiles. When compared with canonical basal markers, NGFR showed a consistent association with survival across both cohorts in gene-by-gene multivariable models. Notably, correlation analyses indicated that NGFR was only weakly correlated with these classical basal markers, whereas the canonical basal markers were more strongly correlated with one another, supporting that NGFR may capture a distinct basal-associated component rather than simply mirroring the expression of the core basal keratin program. These findings position NGFR as a promising marker of a basal-like and prognostically adverse phenotype in BLCA. Nevertheless, although our data is promising, further validation in additional BLCA loss-of-function models and *in vivo* systems will be essential to establish the functional relevance of NGFR and to define its therapeutic potential in BLCA.

In melanoma, the immunoevasive properties of NGFR-expressing cells have boosted the research regarding its roles regulating antitumor cell responses and resistance to ICIs, [25]. Functional studies identified downregulation of antigens and immune ligands regulated by NGFR. The stem-like characteristics of NGFR^+^ tumor cells reduce the number of neoantigens associated with epithelial-differentiated cells, facilitating the evasion of cytotoxic CD8^+^ T cells [24]. Notably, this effect was reversed through pharmacological inhibition of NGFR, reinforcing its driver role. Additionally, NGFR confers resistance to innate immune surveillance by NK cells by reducing the expression of NK-activating ligands [23]. Knockdown of NGFR effectively reversed resistance. In the context of BLCA, NGFR has recently been proposed, through machine learning, as a potential regulator of macrophage polarization [46], suggesting that it may play an important role in immune regulation. We have observed that NGFR is associated with poor response to ICIs in MIBC patients from the IMvigor210 cohort. Additionally, we demonstrate that NGFR correlates with reduced TMB and neoantigens, aligning with the current knowledge in melanoma.

It is difficult to separate prognostic from predictive effects when both can impact survival outcomes. Therefore, our results should be viewed as initial evidence that NGFR is associated with poorer prognosis. Nevertheless, comparisons between the TCGA and IMvigor210 cohorts should be interpreted with caution, as the two cohorts are very different in clinical context: TCGA-BLCA represents a heterogeneous MIBC dataset and IMvigor210 is enriched in locally advanced/metastatic patients treated with atezolizumab. The multivariable models also differed according to the clinical variables available in each cohort; notably, patient age was not available in the downloaded IMvigor210 clinical table, whereas baseline ECOG score was available and was incorporated into that model. These differences likely contribute to the lack of a strictly proportional or identical ordering across all NGFR quartiles, even though both cohorts consistently identify the lowest quartile of NGFR expression as the group with the most favorable survival. An additional limitation of our study is that the clinical associations were derived from transcriptomic datasets, and NGFR mRNA levels may not fully recapitulate protein abundance, localization, or receptor activity in tumor tissues. To address this point, we explored the TCGA-BLCA RPPA dataset; however, NGFR was not represented in the RPPA panel, precluding a direct protein-level validation in this cohort. Therefore, our conclusions should be interpreted at the transcriptomic level, and future studies incorporating NGFR protein-level assessment in clinically annotated BLCA cohorts will be necessary to define its prognostic and predictive value more firmly.

Even in melanoma—where NGFR has been most studied in the context of immune evasion—there are no prospective patient studies confirming a predictive role for ICI resistance. Dedicated prospective cohorts and functional experiments will be needed to determine whether NGFR truly predicts immunotherapy resistance in bladder cancer and other tumors.

To further understand the role of NGFR, we utilized Ecotyper to analyze the tumor ecosystems [34]. Our analysis revealed significant enrichment of stromal components, particularly in CAFs, aligning with an immunosuppressive TME. The contribution of CAFs to the regulation of the antitumor immune response has been studied. CAFs are crucial in shaping the tumor niche, including their interaction with CSCs [17, 49]. In BLCA, by TGFβ signaling CAFs drive resistance to ICIs contributing to a dense stromal compartment that obstructs the effective infiltration of cytotoxic CD8^+^ T cells [32]. Targeting TGFβ in BLCA has demonstrated promising therapeutic success [32]. Our study revealed that NGFR-expressing tumors are particularly enriched in CAFs characterized by markers such as VIM and THY1, as well as genes involved in ECM regulation, TGFβ ligands, and receptors [38]. Interestingly, despite a high CD8 effector signature, NGFR-high showed an immune-excluded phenotype. In these cases, despite the abundance of CD8^+^ T cells, they are confined to the tumor periphery and remain ineffective. These results suggest that, in our cohorts, high NGFR expression is linked to an immune-excluded microenvironment in which CD8+ effector cells are present but remain excluded from the tumor core, likely due to stromal and TGFβ-associated barriers. These results underscore that NGFR-high tumors are associated with an immunosuppressive, immune-excluded TME. Indeed, high NGFR expression is associated with resistance to ICIs in other tumor types [24] correlated with elevated PD-L1 levels [25] or regulating anti-tumor immunity mediated by T and NK cells [23, 24]. Interestingly, our data indicate that NGFR signaling in tumor cells is likely driven by CAF-derived neurotrophins (NGF, BDNF, and NTF3) through paracrine crosstalk, consistent with the enrichment of NGFR-high tumors in stromal-rich carcinoma ecotypes. However, this model remains to be tested experimentally, ideally by perturbing CAF-derived neurotrophins and assessing downstream NGFR activation and tumor cell phenotypes.

Importantly, the immune-excluded phenotype was verified in silico by combining the spatial immune annotation available in IMvigor210 [32] with Ecotyper-derived tumor ecosystems [34] and TGFβ/CAF activity signature [32]. The spatial correlation between NGFR expression and T-cell exclusion described in melanoma [24, 25], suggest a potentially conserved mechanism across cancers. These findings are consistent with NGFR-high tumors being associated with an immune-excluded microenvironment in BLCA, although this will require further validation in independent cohorts with spatially resolved immune profiling.

Therapeutically, NGFR has been targeted using different strategies. Small molecules or a short β-amyloid-derived agonist peptide trigger cell death and slow melanoma tumor growth were used [47, 48]. Similarly, anti-NGFR antibodies have demonstrated success as monotherapies targeting CSCs [49], exhibiting both anti-tumor and anti-metastatic effects. Moreover, strategies that reduce NGFR expression indirectly, such as using heat shock protein inhibitor HSP90 [24] or Ranolazine [50], have led to enhanced anti-tumor immune responses. In addition, we showed that treatment with the NGFR small molecule inhibitor THX-B reduced melanoma lymph node metastasis [45], highlighting its potential as a therapeutic target.

Our study identifies NGFR as a marker of a basal-like cell state in BLCA associated with poor prognosis. In addition, NGFR-high tumors were linked to stromal and immune features consistent with immune exclusion. Although the relationship with immunotherapy outcome requires further validation, these findings suggest that NGFR may help identify a biologically aggressive tumor state with potential clinical relevance. Overall, our results support further investigation of NGFR as a biomarker in BLCA and provide a rationale for future studies exploring whether NGFR-targeting strategies, alone or in combination with immunotherapy, may have therapeutic value in this disease.

## Acknowledgments

This work was supported by Agencia Estatal de Investigación (PID2020-118558RB-I00, PDC2021-121102-I00). J.G.A. was supported by Fundación Científica de la Asociación Española Contra el Cáncer (Predoctoral AECC 2021 PRDMA21602GARC). Work in the lab of F.X.R. was supported, in part, by the Fundación Científica de la Asociación Española Contra el Cáncer (grant PRYGN223005REAL) and the Ministerio de Ciencia e Innovación (grant RTC-2017-6123-1). C.B. was supported by a training grant from the European Union’s Horizon 2020 Research and Innovation Programme under the Marie Sklodowska-Curie grant agreement no. 860895 (TranSYS). M.R. was supported by a Ph.D. fellowship from La Caixa Foundation (LCF/BQ/DR20/11790014).

## Translational relevance

Immunotherapy based on immune checkpoint inhibitors (ICIs) has transformed cancer treatment, outperforming conventional therapies in several tumor types. In bladder cancer (BLCA), ICIs represent a standard treatment for recurrent/metastatic patients. However, resistance is frequent, underscoring the need for biomarkers to personalize treatment. Here, we identify nerve growth factor receptor (NGFR) as a biomarker associated with a lower response probability to ICIs in BLCA. We show NGFR expression in a subset of basal tumor cells and that its high expression is associated with poor prognosis in BLCA. Functionally, we found that NGFR is associated with stemness and invasive capacity in bladder cancer. Furthermore, our data suggest NGFR may act through tumor microenvironment remodeling and activation of cancer-associated fibroblasts (CAFs). Together, these findings position NGFR as a candidate marker of poor prognosis in BLCA, with a potential association with ICI treatment outcome that warrants further clinical validation.

## Supplementary Material

**Supplementary Figure 1.** NGFR marks basal cells in healthy urothelium and BLCA. (**A–D**) Representative images showing immunohistochemistry of NGFR in healthy human urothelium. Scale bar: 100 µm. Sample size: n = 4. (**E**) Representative serial sections of porcine urothelium stained for NGFR and KRT5, showing co-localization of NGFR with basal keratins in the same region. Scale bar: 50 µm. (**F**) Normalized NGFR expression in bladder cancer (BLCA) cell lines from the CCLE consortium, classified as luminal (n = 11) or basal (n = 18). (**G**) Normalized NGFR expression in patient-derived bladder cancer organoids, also classified as luminal or basal [30]. P values for (**F**) and (**G**) were calculated using an unpaired t-test. (**H, I**) GSEA enrichment plots derived from pathway analyses of genes correlated with NGFR expression in basal/squamous (Ba/Sq) CCLE cell lines (n = 18). Shown are the pathways “REACTOME_KERATINIZATION” (NES = 2.62, FDR < 0.001) in (**H**) and “REACTOME_DIFFERENTIATION_OF_KERATINOCYTES_IN_INTERFOLLICU LAR_EPITHELIUUM” (NES = 2.22, FDR < 0.001) in (**I**). P values were computed using permutation-based testing with FDR correction. (**J**) Correlation of NGFR with selected marker genes, including basal markers (red bars, left), epithelial–mesenchymal transition (EMT) markers (yellow bars, middle), and invasion-related markers (blue bars, right).

**Supplementary Figure 2.** NGFR is associated with a BLCA cancer cell population with stem-like and tumorigenic properties. (**A**) Density plots showing NGFR expression levels in the SCaBER CRISPR (CTL), CRISPR NGFR1 (KO1), and CRISPR NGFR2 (KO2) models. (**B, C**) Primary tumorspheres generated in ultra-low-attachment plates from SCaBER CTL, KO1, and KO2 cells. Representative images of the spheroids in CTL or KO tumor cells, and their segmentation, are shown in **B**. The quantification of the total number of tumorspheres is shown in **C**. P-values were calculated using the non-parametric one-way Friedman ANOVA, as the data were not normally distributed. Sample size: n = 4 experiments, with each experiment including three wells run by triplicate per condition. (**D, E**) Transwell migration assays of SCaBER CTL, KO1, or KO2 cells. A representative image of a transwell with planes X and Z (top up, bottom down) is shown in **D**. Quantification of the percentage of migratory cells relative to the total number of DAPI^+^ cells is shown in **E**. P-values were calculated by mixed-effects analysis using one-way ANOVA. Sample size: n = 3 experiments for CTL and KO2 and n = 2 for KO1 (due to not reaching the minimum number of cells per capture), each performed in duplicate per condition and quantified from four images of different transwell sections. (**F**) Representative IHC images of NGFR expression in basal cells in: healthy murine bladder (left), BBN-treated non-malignant bladder (center), and BBN-treated bladder in a malignant state (right). Scale-bars: 50 µm for original, for amplification 10 µm in left and center panels and 20 µm in right panel. (**G**) Representative image of NGFR expression in organoids derived from BBN-induced murine tumors. Scale-bar: 20 µm. (**H**) Representative image of NGFR expression in xenograft tumors generated after subcutaneous inoculation of BBN-derived 2D cells derived from organoids into Nude/Nude immunodeficient mice. Scale-bars: 100 µm for original and 20 µm for amplification.

**Supplementary Figure 3.** Association of NGFR with canonical basal markers in BLCA survival and correlation analyses. (**A, B**) Multivariate survival analysis by Cox proportional hazards models in the TCGA cohort. Each basal marker was evaluated independently adjusting for clinical covariates (**A**) or included with other basal markers to assess their relative prognostic strength (**B**). (**C**) Spearman correlation matrix among NGFR and canonical basal markers in TCGA. (**D, E**) Equivalent analysis in IMvigor210: each basal marker adjusted for clinical variables (**D**) or evaluated in a combined model with other basal markers (**E**). P-values were calculated by Wald test. (**F**) Spearman correlation matrix among NGFR and canonical basal markers in IMvigor210. Basal markers were selected based on their canonical use as markers of the basal program in BLCA. P-values in Cox models were calculated by Wald test.

**Supplementary Figure 4.** NGFR expression is associated with a poor prognostic phenotype and a higher immune and stromal compartment in NMIBC. (**A**) Percentage distribution of NMIBC molecular subtypes [3] as a function of NGFR expression quartiles in the UROMOL cohort. P-values were calculated by Chi-square test (and Fisher’s exact test), comparing the number of patients in each group (corresponding percentage is represented). (**B-D**) Enrichment analysis by ssGSEA of different immune and stromal signatures in the UROMOL2 cohort. Shown are: immune-score signature (**B**), CD8^+^ T-cell deconvolution signature (**C**), and TGF-β signaling signature in fibroblasts (CAFs) (**D**). P-values for comparison between quartiles of NGFR were calculated by one-way ANOVA (ordinary one-way ANOVA).

**Supplementary Figure 5.** Correlation between NGFR expression and its ligands in vitro models, tumor cohorts, and single-cell transcriptomics data. (**A**) Correlation analyses between NGFR and its ligands in basal-subtype bladder cancer cell lines from the CCLE dataset (n = 18). Correlations between NGFR and all neurotrophin ligands is shown. (**B**) Correlation between NGFR and its ligands in primary MIBC tumors from the TCGA-BLCA cohort (bulk RNA-seq). (**C**) Correlation between NGFR and its ligands in MIBC tumors from the IMvigor210 cohort (bulk RNA-seq). The p-values were computed using Pearson’s r correlation analysis. Both p-values and Pearson’s r correlation value are indicated in panels (**A**), (**B**) and (**C**). (**D, E**) Single-cell RNA-seq analysis of NGFR ligand expression across cell types. A dot plot (**D**) and UMAP projection (**E**) illustrating the distribution of ligand expression are shown. See Figure 1E for scRNA-seq cell type annotations. (**F-I**) Correlation analyses in the IMvigor210 cohort between NGFR expression and TMB (**F**), neoantigen burden (**G**), CD8^+^ effector signature (**H**), and TGF-β signaling signature in CAFs (**I**). The p-values were computed using Pearson’s r correlation analysis, and both p-values and Pearson’s r correlation value are indicated in each panel.

**Supplementary Figure 6.** Analysis of the relevance of NGFR in the cohort of patients MBIC from the IMvigor010 study. (**A, B**) Survival analysis in the IMvigor010 cohort. Kaplan–Meier curves for the entire cohort (**A**) and the subgroup of patients treated with atezolizumab (**B**), categorized into quartiles according to NGFR expression (Q1: lowest expression; Q4: highest expression). Disease-free survival (DFS) is shown; p values were calculated using the log-rank test. The number of patients per quartile: n = 182 per quartile in the entire cohort and Q1 = 85, Q2 = 97, Q3 = 96, Q4 = 96 in the atezolizumab-treated group. (**C**) Distribution of patients with or without relapse after ICI treatment in the atezolizumab arm, stratified by NGFR expression quartiles. Error bars indicate 95% binomial confidence intervals for the proportion of patients with relapse in each quartile. The p value corresponds to the overall comparison of relapse proportions across quartiles using the chi-square test. (**D**) Distribution of consensus molecular subtypes [6] based on NGFR expression quartiles in the IMvigor010 cohort. P values were calculated using the chi-square test, and the relative percentages of each subtype per quartile are shown. (**E**) NGFR expression in patients classified as TMB^low^ and TMB^high^, according to the clinical annotation of the cohort (n = 382 and n = 191, respectively). The p-value was calculated using the nonparametric Mann–Whitney test. (**F**) Association between NGFR expression (by quartiles) and TGFβ signature in cancer-associated fibroblasts (CAFs). The p-value was obtained using one-way ANOVA.

