## Supplementary figures and images for "Nerve growth factor receptor identifies a basal subpopulation linked to poor prognosis and reduced immunotherapy responses in bladder cancer"

### Supplementary Figure 1

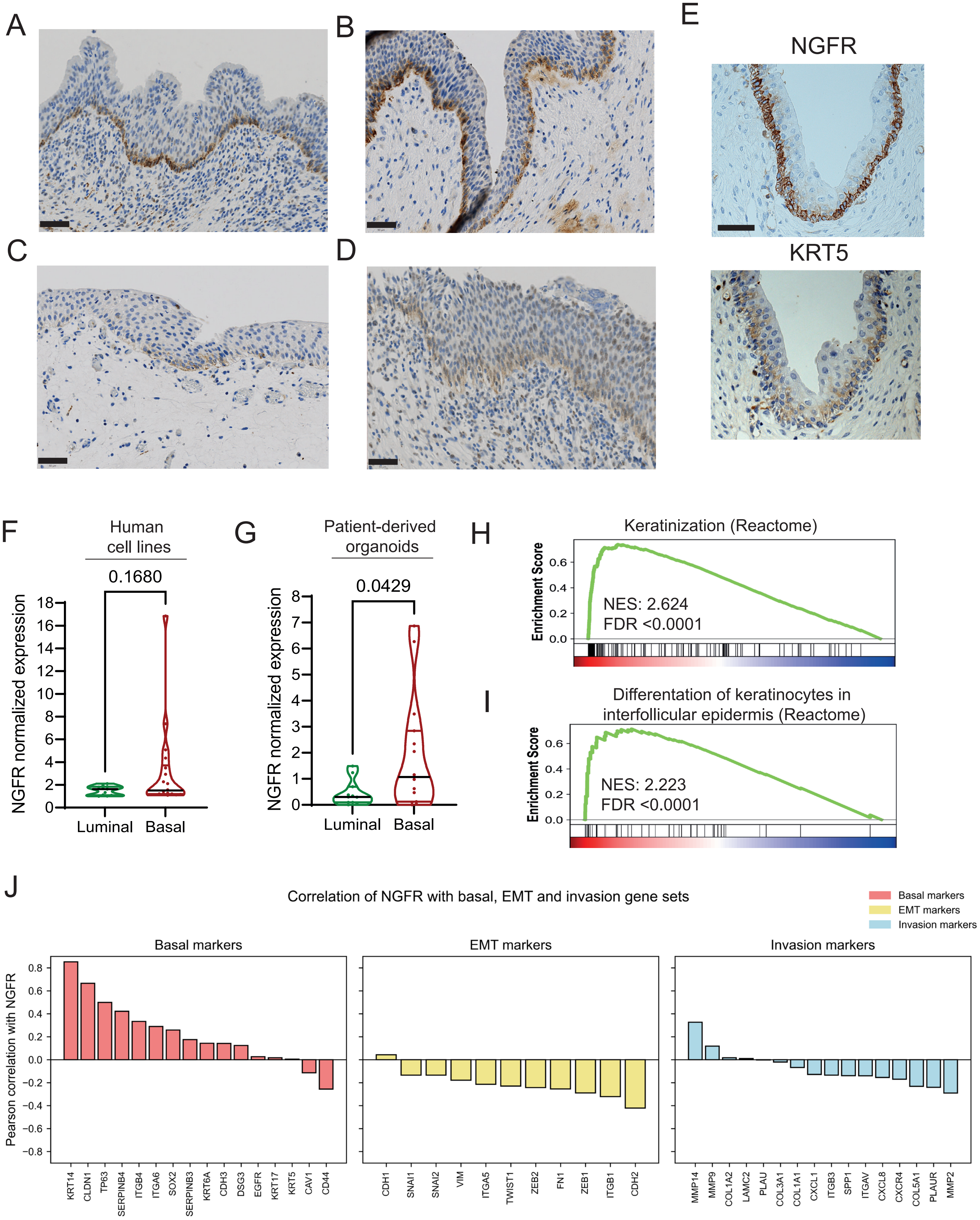


#### Supplementary Figure 1

### Supplementary Figure 2

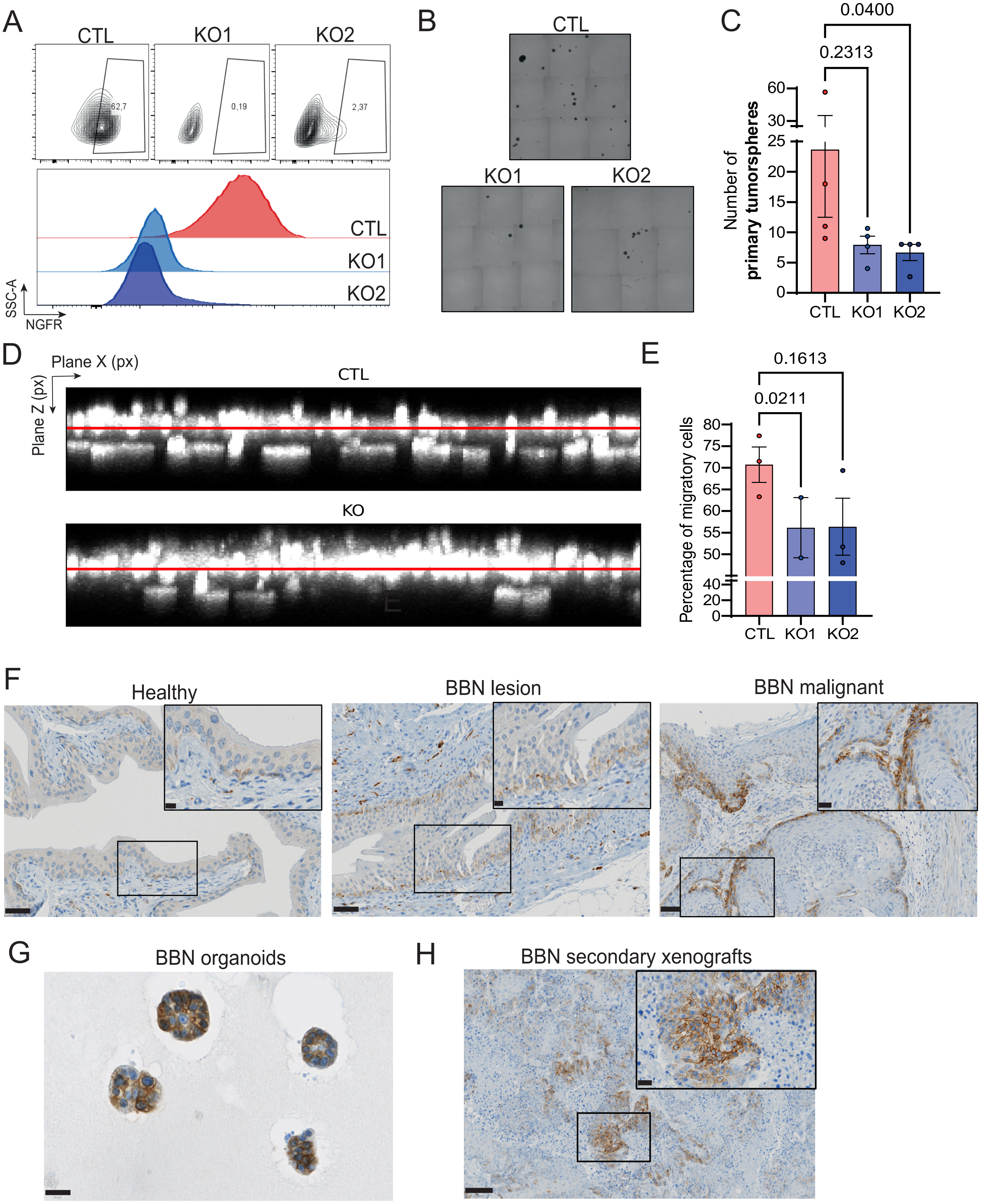


#### Supplementary Figure 2

### Supplementary Figure 4

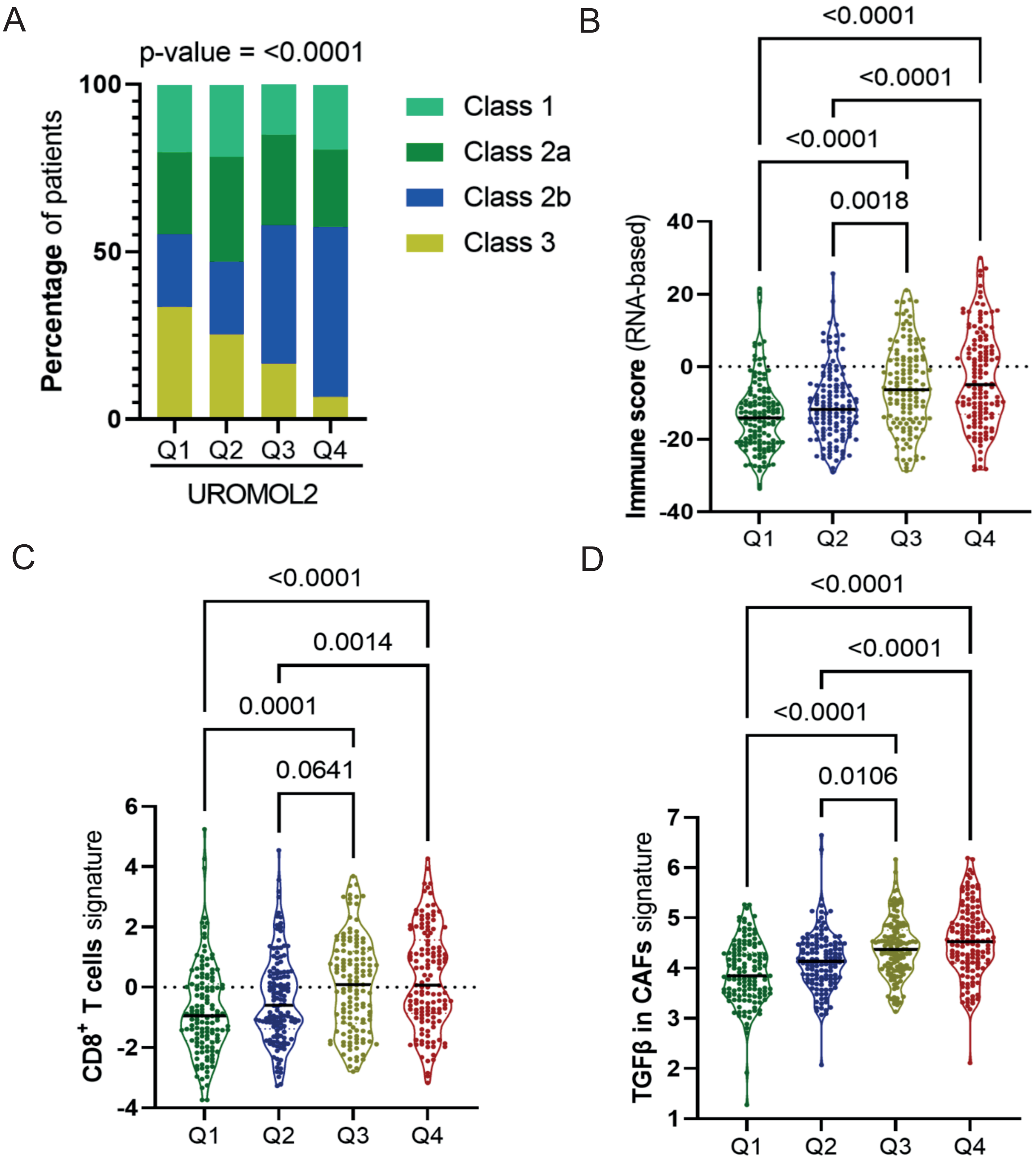


Supplementary Figure 4

### Supplementary Figure 5

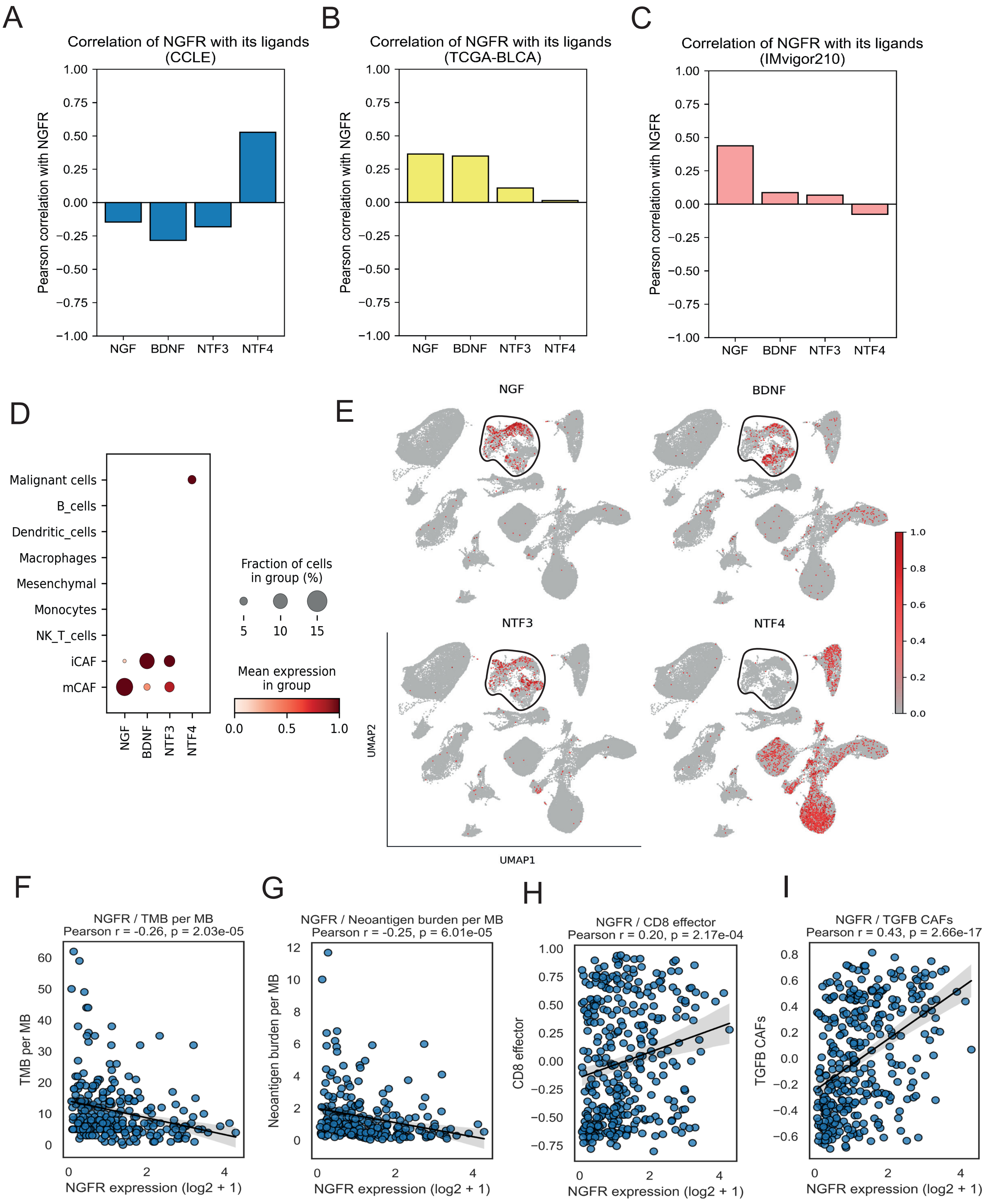


#### Supplementary Figure 5

### Supplementary Figure 6

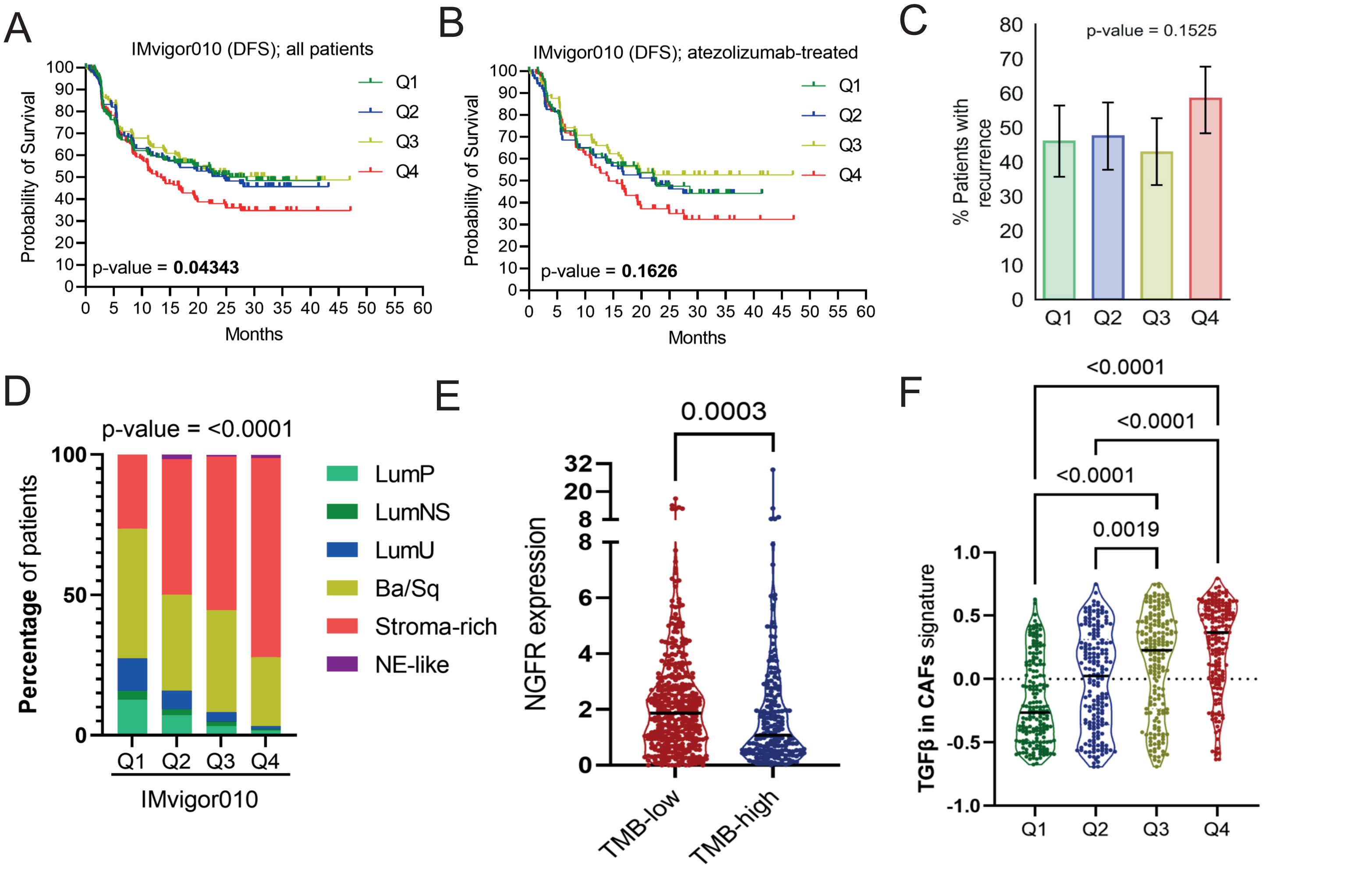


Supplementary Figure 6
