## Supplementary Figure 3 for "Nerve growth factor receptor identifies a basal subpopulation linked to poor prognosis and reduced immunotherapy responses in bladder cancer"

A

| Basal Genes | HR | Lower CI | Upper CI | p |
|---|---|---|---|---|
| **NGFR** | 1.219 | 0.044 | 0.352 | 0.012 |
| **KRT5** | 1.242 | 0.057 | 0.377 | 0.008 |
| **KRT14** | 1.156 | -0.017 | 0.308 | 0.079 |
| **KRT6B** | 1.209 | 0.036 | 0.344 | 0.016 |
| **CD44** | 1.112 | -0.058 | 0.270 | 0.204 |

B

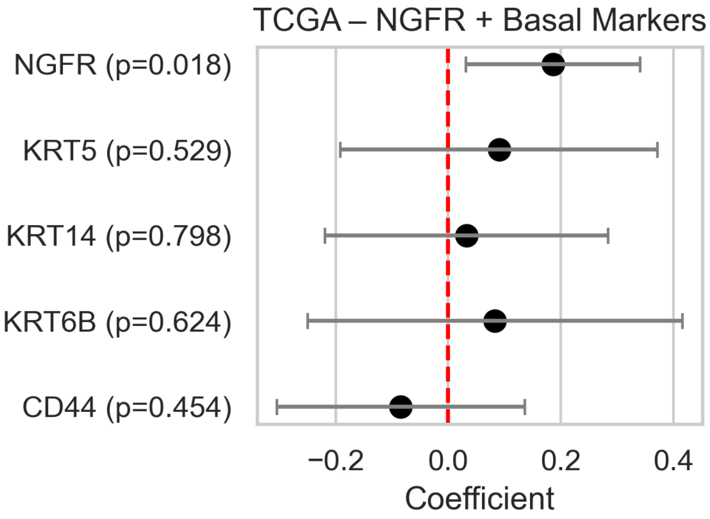


C

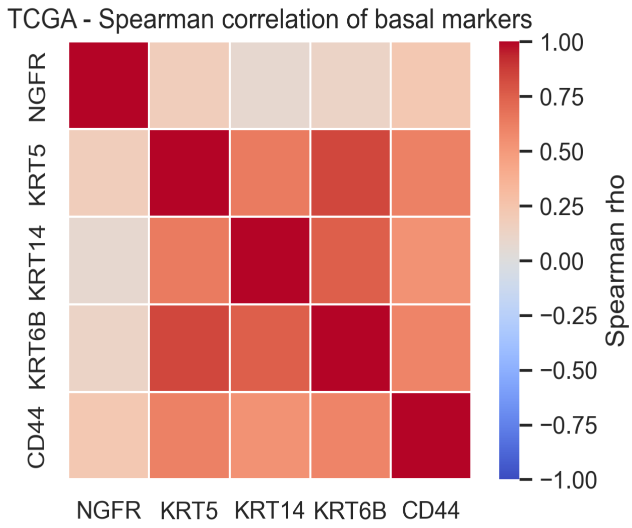


D

| Basal Genes | HR | Lower CI | Upper CI | p |
|---|---|---|---|---|
| **NGFR** | 1.206 | 0.041 | 0.335 | 0.012 |
| **KRT5** | 1.049 | 0.007 | 0.089 | 0.023 |
| **KRT14** | 1.023 | -0.017 | 0.063 | 0.251 |
| **KRT6B** | 1.046 | 0.002 | 0.088 | 0.039 |
| **CD44** | 1.111 | -0.006 | 0.216 | 0.064 |

E

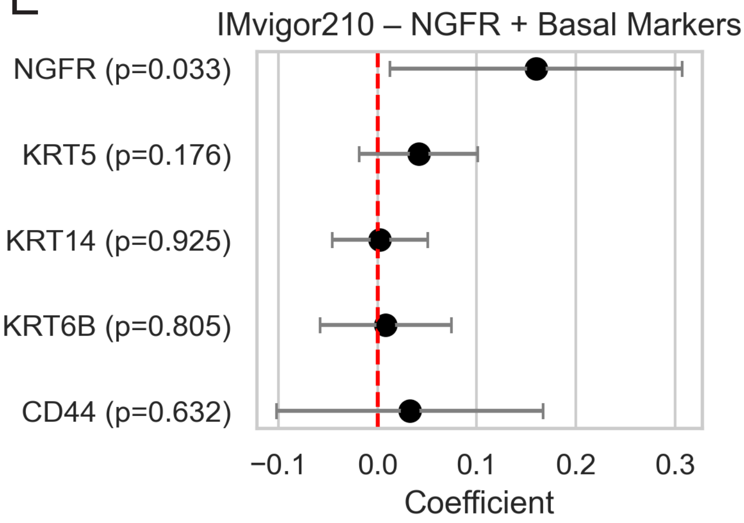


F

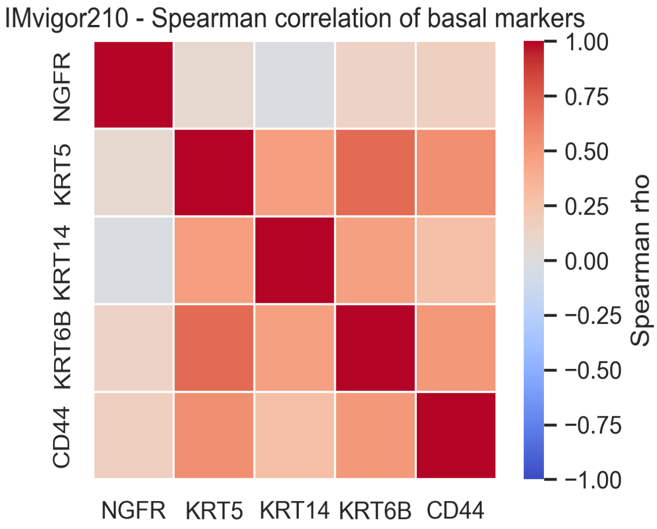


### Supplementary Figure 3
